# Efficiency Improvements and Discovery of New Substrates for a SARS-CoV-2 Main Protease FRET Assay

**DOI:** 10.1101/2021.02.19.431973

**Authors:** Tonko Dražić, Nikos Kühl, Mila M. Leuthold, Mira A. M. Behnam, Christian D. Klein

## Abstract

The COVID-19 pandemic, caused by the SARS-CoV-2 virus, has a huge impact on the world. Although several vaccines have recently reached the market, the development of specific antiviral drugs against SARS-CoV-2 is an important additional strategy in fighting the pandemic. One of the most promising pharmacological targets is the viral main protease (M^pro^). Here, we present an optimized biochemical assay procedure for SARS-CoV-2 M^pro^. We have comprehensively investigated the influence of different buffer components and conditions on the assay performance, and characterized six FRET substrates with a 2-Abz/Tyr(3-NO_2_) FRET pair. The substrates 2-AbzSAVLQSGTyr(3-NO_2_)R-OH, a truncated version of the established DABCYL/EDANS FRET substrate, and a new substrate 2-AbzVVTLQSGTyr(3-NO_2_)R-OH are promising candidates for screening and inhibitor characterization. In the latter substrate, the incorporation of Val at the position P5 improved the catalytic efficacy. Based on the obtained results, we present here a reproducible, reliable assay protocol using highly affordable buffer components.

## INTRODUCTION

The coronaviridae are a family of positive-strand RNA viruses which infect mammals, birds, reptiles and amphibians. In humans, several types of coronavirus are important as pathogens causing a plethora of illness ranging from mild respiratory infections to the life threatening disease such as severe acute respiratory syndrome (SARS). Coronaviruses are the pathogens that led to the SARS pandemic in 2002/2003, the MERS epidemic (ongoing since 2012) and the present COVID-19 pandemic (from 2019).^1–3^ The viral disease COVID-19 is caused by SARS-CoV-2, which was first described in Wuhan in December 2019.^4^ Today the virus has spread to many countries and turned into a worldwide pandemic with currently (as of January 25, 2021) more than 99 million infected people. The death toll associated with the virus is more than 2.1 million cases.^5^ In around 80% of the symptomatic infections, a mild disease with fever or mild pneumonia can be observed, in 14% of the cases it is more severe and about 5% patients have to be treated in intensive care units.^6^ The infection usually occurs through a close contact and aerosols in social interaction.^7^

Although the first vaccines have already been approved, there is a lack of effective drugs against the viral infection. Since the viral main protease M^pro^ (3CL^pro^, nsp5) has no closely related homologues in humans and is involved in an essential step of the viral life cycle, it is one the most promising drug targets.^8^ The chymotrypsin-like cysteine protease M^pro^ cleaves the viral polyprotein into several functional proteins. For efficient proteolytic activity, it requires dimerization with a second protease molecule. It cleaves the polyproteins at 11 distinct sites, exclusively after glutamine residues.^8–10^ Many main protease inhibitors have been described, most having a peptidic structure and containing a reactive electrophilic group for covalent binding to the catalytic cysteine.^11–16^

Biochemical assays are valuable tools to study viral proteases and to develop and improve protease inhibitors. A frequently used and high-throughput capable technique to investigate the activity of proteases are Forster resonance energy transfer (FRET)-based assays. In FRET, the energy of an excited donor functionality is transferred to an acceptor moiety in close proximity, the “quencher”. The energy transfer is radiation-free and therefore not exchanged via emission and absorption of photons. By incorporation of a FRET pair into an enzyme substrate, the leavage rate of the substrate can be investigated.^17–18^

We herein present the development of a biochemical FRET-based SARS-CoV-2 M^pro^ assay. Several assay conditions and additives such as salts, polyols and detergents were studied. Furthermore, different established and new FRET substrates were synthesized, compared and tested.

## RESULTS AND DISCUSSION

### Assay conditions

In order to find the optimal assay conditions, measurements with substrate **4** (see below) and various assay conditions as well as buffer components were carried out (Figure 1). First, the influence of different buffers was investigated. Most SARS-CoV-2 M^pro^ biochemical assays were conducted using Tris buffer^9, 12–15, 19–21^ or FIEPES buffer.^11, 22–23^ We found no significant difference between Tris and phosphate buffer, whereas the fluorescence increase over time was lower when using a HEPES buffer (Figure 1A). Next, the addition of different salt concentrations was examined. The ionic strength had no obvious influence on substrate cleavability. This is in contrast to a previously published study on SARS-CoV M^pro^.^24^ Other additives like the reducing agents dithiothreitol (DTT) and tris(2-carboxyethyl)phosphine (TCEP) led to higher enzyme activity (Figure IB). Reducing agents are often included in buffers for enzyme storage, purification and biochemical assays to prevent cysteine oxidation. However, electrophilic inhibitors that are reactive towards cysteine residues may be scavenged by thiols in the assay buffer, thereby yielding false-negative results. Our results indicate that there is no necessity for thiols in the assay buffer, so that screening protocols may be implemented that allow the search for and characterization of thiol-reactive inhibitors with a covalent binding mode to the catalytic cysteine.

**Figure 1.**
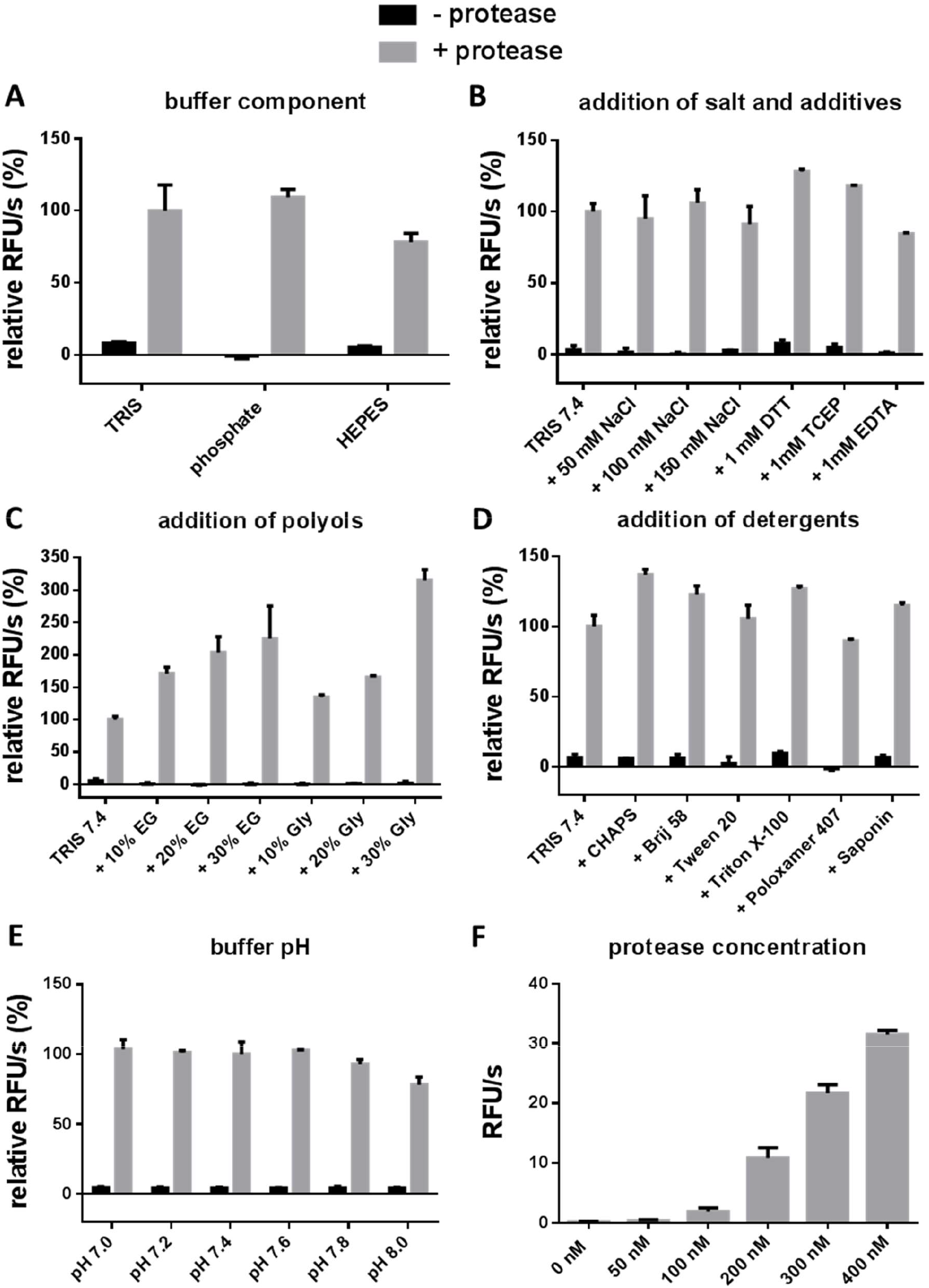
Buffer optimization of the FRET assay. The measurements were performed with substrate **4** (50 μM) and 500 nM enzyme concentration and at pH 7.4 unless indicated otherwise. Relative RFU/s are given in comparison to the RFU/s of the cleavage reaction in Tris buffer without additives. (A) Comparison of substrate cleavage velocity with different buffer components, pH 7.5. (B) Influence of salts and additives. (C) Influence of polyols. EG = ethylene glycol, Gly = glycerol. (D) Influence of detergents at a concentration of 0.01%. (E) Influence of pH (Tris buffer). F) Different enzyme concentrations. All measurements were performed in triplicate.

The chelating agent EDTA reduced the enzyme activity slightly. Using polyols as additives, we observed the highest enzymatic activities at high concentrations (Figure 1C). 30% glycerol led to a higher substrate cleavage compared to 30% ethylene glycol but also caused a high viscosity of the assay buffer which interfered with pipetting. We also studied the effect of ionic and nonionic detergents on enzyme activity. All detergents were assayed at a concentration of 0.01%. Inclusion of detergents into the assay buffer can prevent the formation of colloidal aggregates that lead to non-specific inhibition.^25–27^ All detergents except poloxamer 407 increased the enzyme activity (Figure ID). The highest increase was measured with the zwitterionic detergent CHAPS. However, it should be considered that ionic functionalities may interact with charged functionalities of the test compounds leading to false-negative results.

Furthermore, we tested the influence of different pH values on substrate cleavage velocity and found no significant influence in the pH range from 7.0 to 8.0 (Figure 1E). A minimal decrease in activity can only be seen at a basic pH value higher than 7.6. Several publications described the influence of pH on enzyme activity of the SARS-CoV M^pro^. Fan et al. and Tan et al. reported a peak of substrate cleavage at pH 7.0^28–29^, whereas other studies reported the highest processing at around pH 7.5^30^ or pH 8.O.^24,31^ Since the pH influences the conformation of the protease^28^ and the inhibitor recognition, a physiological pH value should be chosen. Additionally, various protease concentrations and the C144A mutant were investigated. With increasing enzyme concentrations, the substrate cleavage increased significantly (Figure 1F). In order to avoid reaching the assay wall in compound screenings while maintaining high signal intensity, a moderate enzyme concentration of 300 nM was selected. The mutant protease proved to be inactive (Figure SI).

For further investigations, we used the combination of 50 mM Tris-HCl pH 7.4, 100 mM NaCl, 20% ethylene glycol and 0.0016% Brij 58 because this ensures a high activity of the protease as well as reliability and easy handling.

### Substrate characterization

After establishing the assay conditions, we proceeded with the investigation of the substrates. Eight FRET substrates were designed and synthesized, with *N*-terminal 2-aminobenzoic acid as a fluorophore (Figure 2A). As a C-terminal quencher either 3-nitrotyrosine (Tyr(3-NO_2_)-OH; substrates **1, 2, 3, 5,** and **6**) or *N*-beta-(2,4-dinitrophenyl)-L-2,3-diaminopropionic acid (L-Dap(Dnp)-OH; substrate **4**) was used. The substrates were synthesized by solid phase peptide synthesis (SPPS) using the Fmoc strategy. The amino acid sequences were chosen based on the cleavage preferences of M^pro^ and substrates developed for SARS-CoV M^pro^ biochemical assays. Substrate **1** contains a truncated sequence from the FRET substrate DABCYL-KTSAVLQSGFRKME-EDANS, which was established for the assays of SARS-CoV-1 M^pro^ ^29, 32^, but also commonly used in the SARS-CoV-2 assays.^11–12, 14, 22, 33^ Substrate **2** was introduced by Blanchard et al. for the high-throughput screening of inhibitors against SARS-CoV-1 M^pro 34^, while substrate **3** is a serendipitous discovery. In substrate **4**, the quencher 3-nitrotyrosine was replaced by Dap(Dnp)-OH in order to exclude potential fluctuations of fluorescence in the measurement, incurred from the use of an assay buffer with similar pH value to the pKa value of Tyr(3-NO_2_)-OH side chain. Substrate **5** was previously recognized as a sequence not requiring glutamine at P1 position^35^, whereas substrate **6** was designed to test the minimal sequence required for the recognition. Substrates **7** and **8** are substrates developed and routinely used in our group for the dengue and West Nile vims NS2B/NS3 protease assays, and were used here as negative controls.^36–37^

**Figure 2.**
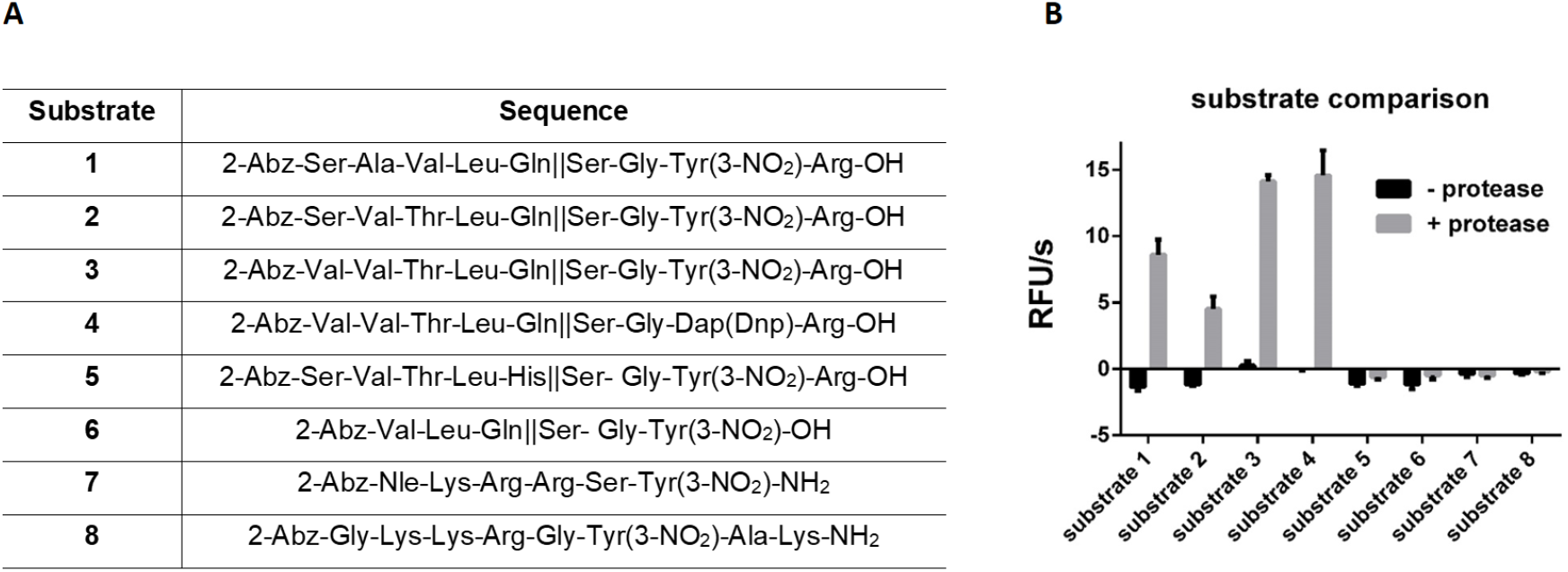
(A) The sequences of the substrates **1-8.** (B) Comparison of cleavage rates of different substrates at 300 nM enzyme concentration. All measurements were performed in triplicate.

We first screened the activity of the protease with substrates at a concentration of 50 uM. The highest cleavage was observed for substrates **3** and **4** (Figure 2B). These two substrates contain the amino acid sequence derived from the SARS-CoV-1 substrate **2**, with a serendipitous replacement of serine for valine at the position P5. This change caused almost 3-fold increase in the cleavage of the substrate. Substrate **1** showed activity in between substrates **2** and **3** and **4**, whereas substrates **5**, with histidine at PI, and the minimal sequence substrate **6** had no activity. As expected, DENY and WNV substrates (**7** and **8**) also exhibited no cleavage.

By determining *K_m_*, *V*_max_, *K*_cat_ and *K*_cat_/*K*_m_ values, we further characterized four substrates that displayed cleavage in the initial screening (Figure 3 and Table 1). In the calculations of *K*_m_ and *V*_max_, we have included a correction for the inner filter effect of FRET pairs for each individual substrate (Tables S1 - S4).^36, 38^ Substrate **3** showed the highest catalytic efficiency (*K*_cat_/*K*_m_) with a value 441.3 M^−1^s^−1^. Substrate **4** with the same amino acid sequence, but different quencher, had an almost 2-fold lower *K*_cat_/*K*_m_ value of 265.3 M^−1^s^−1^. This was in the same range as for substrates **1** and **2**, with *K*_cat_/*K*_m_ 272.2 M^−1^s^−^1 and 241.4 M^−1^s^−1^ respectively. Substrate **4** showed the lowest *K*_m_ value, but also had the lowest *V*_max_, whereas for substrate **1**, both values were the highest of the four characterized substrates. The *K*_cat_/*K*_m_ value for substrate **1** is more than 10-fold lower than the values for the full sequence equivalent substrate (DABCYL-KTSAVLQSGFRKME-EDANS) reported in the literature (*K*_cat_/*K*_m_ ranges from 3011 M^−1^s^−1^ for SARS-CoV-1 and 3426 M^−1^s^−1^ for SARS-CoV-2^12^ to 5624 M^−1^s^−1^ ^11^ and 6689 M^−1^s^−1^ ^22^ reported for SARS-CoV-2). However, the full sequence substrate showed comparable catalytic efficacy (214 M^−1^s^−1^) to our substrate **1**, when measured with the native SARS-CoV-2 M^pro^ with the two extra residues histidine and methionine at the iV-terminus.^22^ Moreover, the truncated substrate TSAVLQSGFRK displayed similar catalytic efficacy (177 M^−1^s^−1^) in combination with the SARS-CoV-1 M^pro^.^29^ The previously reported *K*_cat_/*K*_m_ value for substrate **2** was 20 M^−1^s^−1^ for SARS-CoV-1.^34^ This is 10-fold lower than the value obtained in our assay, however it has to be noted that in the previous work, the assay was performed in phosphate buffer without the addition of other components. This substrate has not been described on SARS-CoV-2 M^pro^ so far.

**Table 1.** *K*_m_, *V*_max_, *K*_cat_ and *K*_cat_/*K*_m_ values for different substrates determined using FRET assay.

| substrate | 1 | 2 | 3 | 4 |
| --- | --- | --- | --- | --- |
| $K_m$ [$\mu\text{M}$] | $375.9 \pm 78.3$ | $100.8 \pm 22.8$ | $132.2 \pm 33.6$ | $77.9 \pm 19.2$ |
| $V_{\text{max}}$ [$\mu\text{M/s}$] | $0.0307 \pm 0.0038$ | $0.0073 \pm 0.0006$ | $0.0175 \pm 0.0018$ | $0.0062 \pm 0.0005$ |
| $K_{\text{cat}}$ [$\text{s}^{-1}$] | 0.1023 | 0.0243 | 0.0583 | 0.0207 |
| $K_{\text{cat}}/K_m$ [$\text{M}^{-1}\text{s}^{-1}$] | 272.2 | 241.4 | 441.3 | 265.3 |

All measurements were carried out in triplicate.

**Figure 3.**
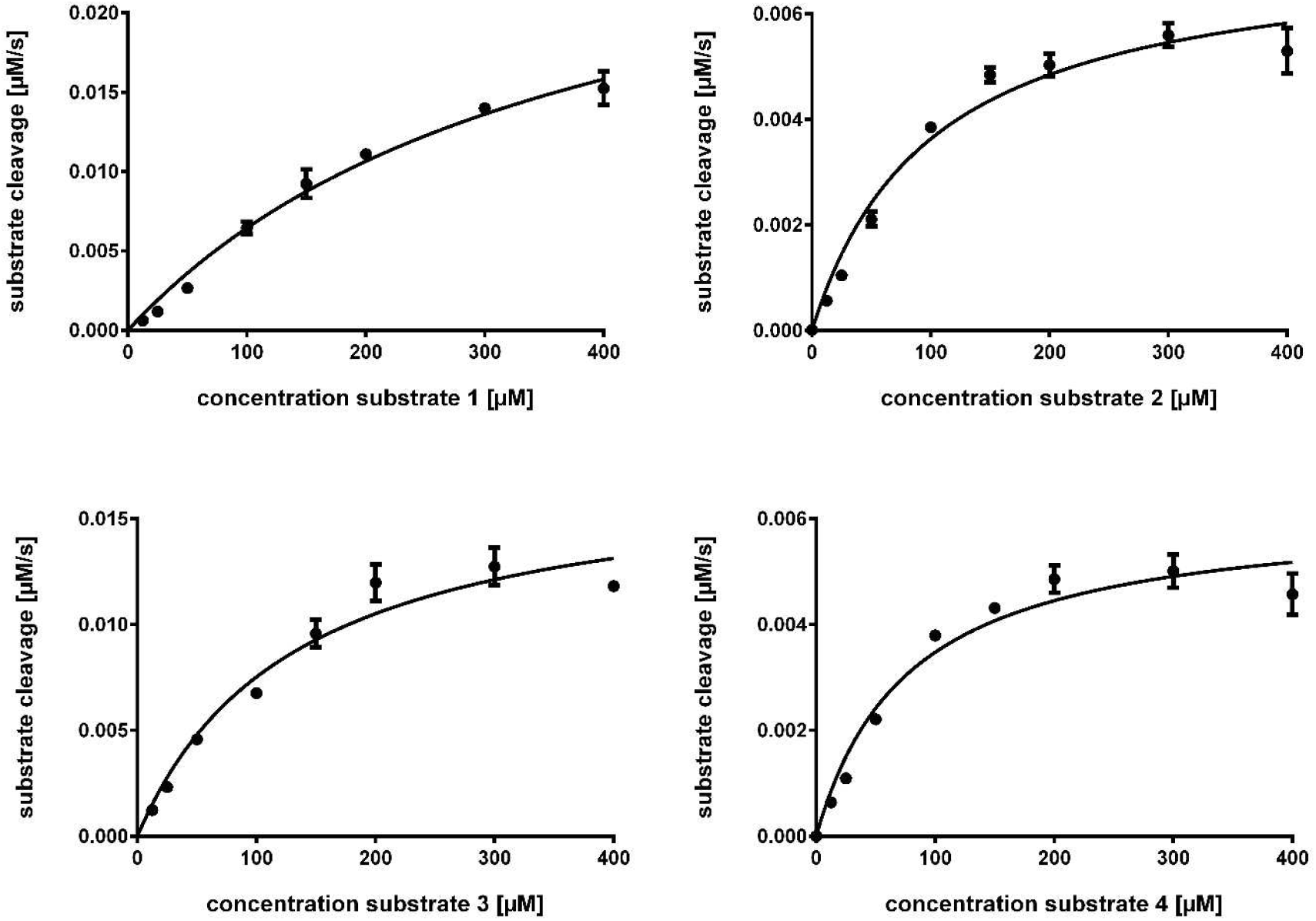
Michaelis-Menten analysis of the interaction of FRET substrates with the SARS-CoV-2 M^pro^. All curves were corrected for inner filter effects.^38^ All measurements performed in triplicate.

### Z’ Score

The Z’-score determination for the assay was performed with substrate **1** at a protease concentration of 300 nM (Figure 4). The calculated score of 0.65 indicates high reproducibility, robustness, and reliability of the assay. Furthermore the score and a signal window greater than 2 demonstrate excellent assay performance and capability for high throughput screenings.^39–40^

**Figure 4.**
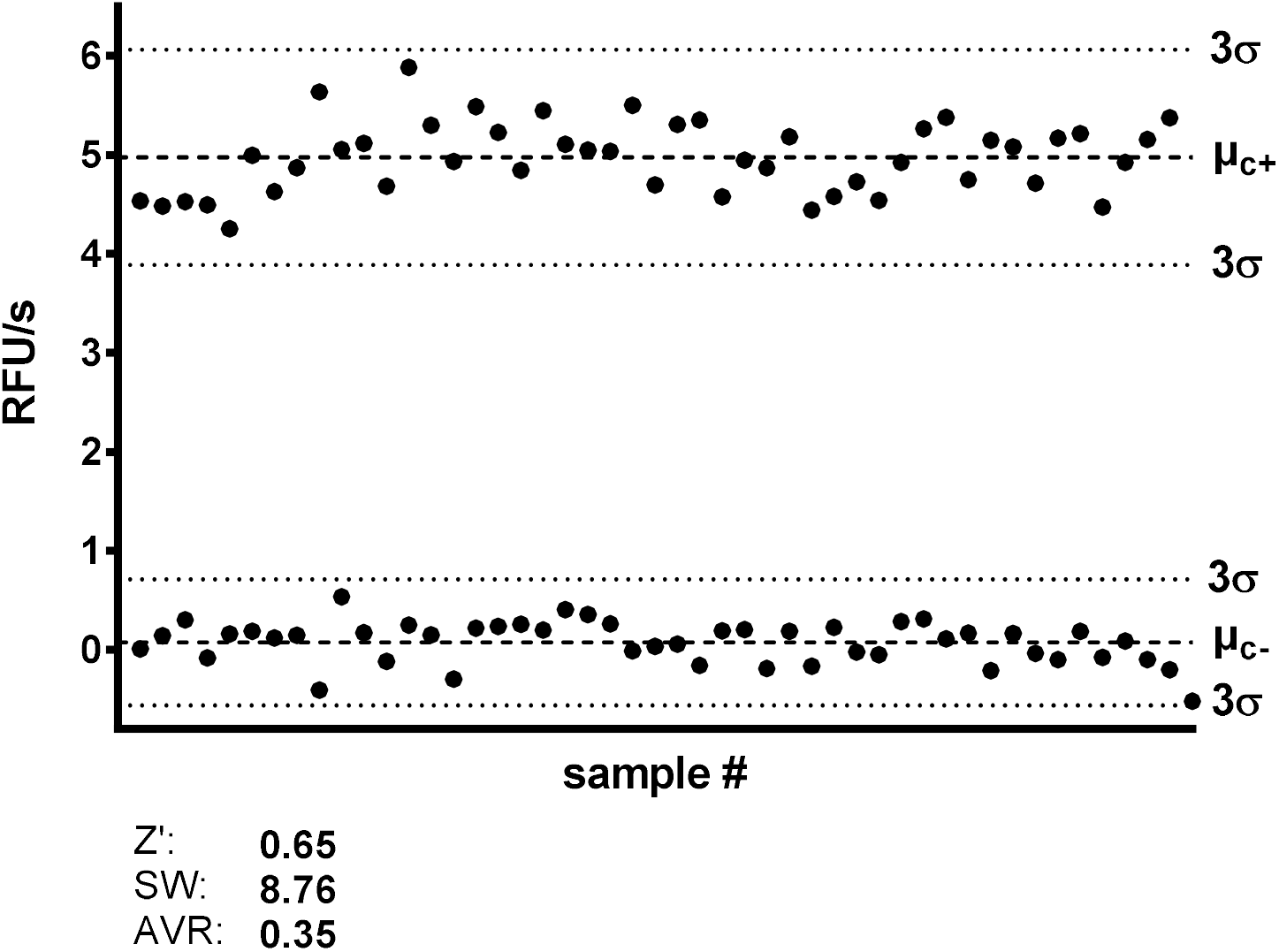
Z’-factor and signal window determination.

### Reference inhibitors

Substrate **1** was chosen for screening and IC_50_ determinations of the reference inhibitors (cf. Figure 5, Table 2). Three known inhibitors were selected as reference compounds according to their previously published activities against SARS-CoV-1 and SARS-CoV-2 M^pro^. The FDA-approved hepatitis C protease inhibitor **boceprevir** was previously described as a potent covalent active-site inhibitor of SARS-CoV-2 M^pro^ with low micromolar IC_50_ values.^11, 15, 41^ The thiophene chloropyridinyl ester **MAC-5576** was identified as a potent SARS-CoV-1 M^pro^ inhibitor in a high-throughput screening campaign.^34^ Several structure-activity studies were earned out which led to improved inhibitory activity of the compound series.^42–44^ The furoic ester **FE-1** was detected as one of the most active SARS-CoV-1 M^pro^ inhibitors. Furthermore it was shown that both esters are active-site-directed covalent inhibitors.^42^ In 2020 this compound series was also described as active against the SARS-CoV-2 M^pro^ with a nanomolar IC_50_ value for compound **MAC-5576**.^45–46^ Since all compounds are active site inhibitors, IC_50_ value differences to the reference values are most likely due to different substrate concentrations and K_m_ values in the cited works. Additionally, IC_50_ values of covalent inhibitors are dependent on assay incubation times which differed from assay to assay.

**Table 2.**
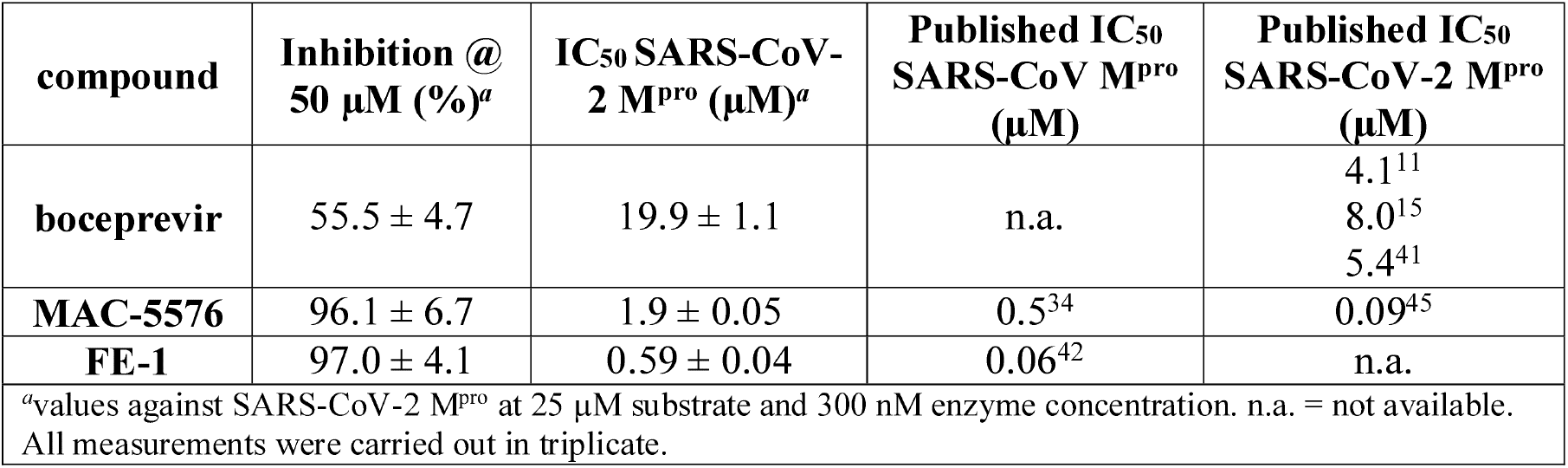
Inhibitory Activity of Reference Compounds against Isolated SARS-CoV-2 M^pro^ and published IC_50_ values against SARS-CoV and SARS-CoV-2 M^pro^.

**Figure 5.**
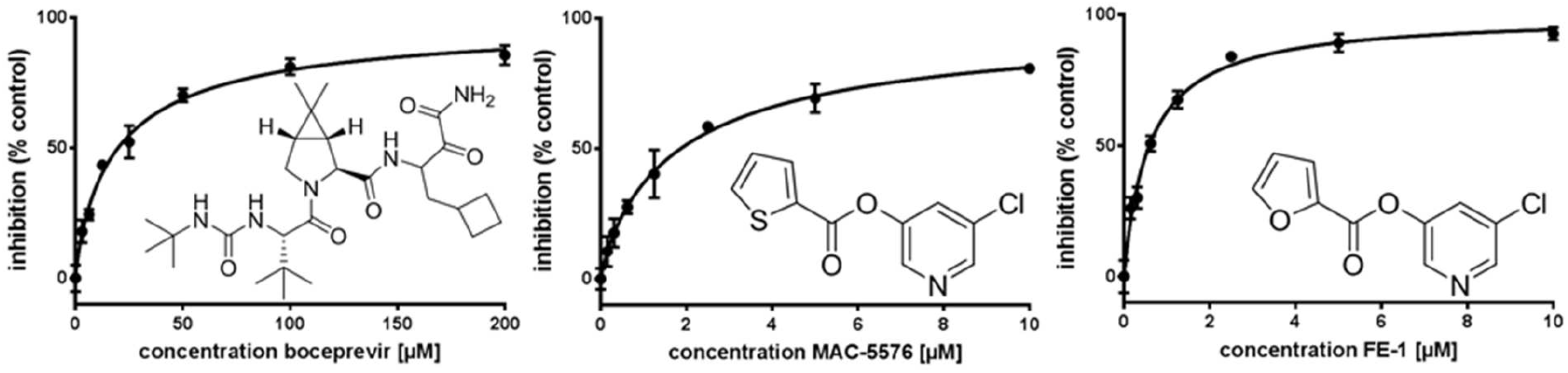
Dose-response curves and structures of reference compounds **boceprevir**, **MAC-5576** and **FE-1.** R square values for all curves are > 0.99. All measurements were performed in triplicate.

## CONCLUSIONS

We have performed a systematic evaluation of assay conditions for the SARS-CoV-2 protease. We have shown that the performance with Tris and phosphate buffer is higher than with HEPES. The addition of salts had no influence on the protease activity, whereas polyols, as well as most of the tested detergents improved activity. We have designed and tested six substrates under the newly established assay conditions. Substrate **1** with the FRET pair 2-Abz/Tyr(3-NO_2_), which is a truncated version of the commonly used DABCYL-KTSAVLQSGFRKME-EDANS substrate, has shown to be reliable and sufficiently active under our assay conditions and is suitable for performing high throughput assays. By decreasing the size by 5 amino acid residues and replacing the DABCYL/EDANS FRET pair for the less expensive 2-Ab//Tyr(3-NO_2_), it is possible to minimize costs and time of the assay preparation, particularly in the preparation of the substrate, while maintaining good performance. Furthermore, we have shown that in substrate **2** a replacement of only one amino acid, *i.e.* serine for valine at the position P5, leads to an improved catalytic efficacy of the substrate. This observation also provides some additional insights to the general substrate - and possibly inhibitor - recognition properties of this enzyme and the main proteases of other coronaviruses with similar active sites. It may be speculated that the small substrate presented here has a higher accessibility to the active site of the MPro under in-vitro conditions than the bulkier FRET substrates described previously: MPro has a tendency to form dimers in solution, which may occlude or restrict access to the active site and interfere with free diffusion of the substrate.

In conclusion, we have developed a reliable and reproducible biochemical assay for the SARS-CoV-2 main protease, which can be applied in high throughput screenings and focused characterization of inhibitors. We hope that the newly discovered conditions and substrates will aid the development of potent antiviral compounds against SARS-CoV-2.

## EXPERIMENTAL SECTION

### General comments

All chemicals were purchased from commercial suppliers. **Boceprevir** was obtained from Biosynth Carbosynt. The measurements were performed in black 96-well V-bottom plates (Greiner Bio-One, Germany) using a BMG Labtech Fluostar OPTIMA Microtiter fluorescence plate reader at an excitation wavelength of 330 nm and an emission wavelength of 430 nm. All measurements were performed at room temperature.

### Construct Design

The expression construct for the SARS-CoV-2 M^pro^ was designed by multiple sequence alignment of Wuhan seafood market pneumonia virus isolates 2019-nCoV (accession numbers MN938384.1; MN975262.1; MN988713.1; MN985325.1). The reading frame for the SARS-CoV-2 M^pro^ was determined by alignment with previous SARS-CoV M^pro^ expression constructs.^28^ Codon usage was optimized individually for optimized expression in both *E. coli* and eukaiyotic systems. The gene sequence coding for the SARS-CoV-2 M^pro^ protease was ordered at Eurofins with Notl and BamHI restriction sites at the 3’ and 5’ ends, respectively.

The gene sequence encoding the SARS-CoV-2 M^pro^ was inserted by restriction-based cloning into a pET28a(+) expression vector in a way to have a C-terminal His-tag, since structure analysis of the SARS-CoV-1 protease indicated less disruption of dimerization using a C-terminal tag compared to an N-terminal tag. The cleavage inactive mutant C144A was cloned from the wild-type construct by molecular assembly with two sets of primers. Identity of the constructs was determined by agarose gel electrophoresis, colony PCR, and sequencing.

### Expression and Purification of SARS-CoV-2 M^pro^

Plasmid encoding the SARS-CoV-2 M^pro^ was transformed into *E. coli* BL21 DE3 cells for expression. Overnight culture from a single colony was grown shaking at 37 °C in LB medium supplemented with 50 mg/ml kanamycin. The next day, pre-warmed LB medium with kanamycin was mixed 1:40 with overnight culture and bacteria were grown until optical density at 600 nm reached 0.3 - 0.4. Subsequently, temperature was reduced to 25 °C. Protein expression was induced by addition of 1 mM isopropyl-D-thiogalactoside (IPTG) and cells further grown shaking for 24 h at 25 °C. Bacteria were harvested by centrifugation, the resulting cell pellet flash frozen in liquid nitrogen and stored for further use at −80 °C.

Purification of the SARS-CoV-2 M^pro^ was performed on ice or at 4 °C. 1 g cell pellet was thawed in 10 ml ice-cold buffer A (20 mM Tris, 200 mM NaCl, 10 mM imidazole, pH 7.6). Dissolved bacteria were lysed using a cell disrupter (Constant Systems LTD.). Cell debris was centrifuged 2 h at 50’000 x g, to remove insoluble aggregates and inclusion bodies. Supernatant after centrifugation was mixed with 1 ml pre-equilibrated (buffer A) nickel-beads and incubated on a rolling shaker for 30 min at 4 °C. Subsequently nickel beads were washed with each 1020 ml buffer A with increasing concentrations of imidazole (10 mM, 20 mM, 50 mM, pH 7.6). Protein was eluted by multiple elution steps using each 1ml of buffer B (20 mM Tris, 200 mM NaCl, 500 mM imidazole). Elution fractions were analyzed by OD measurement and SDS-PAGE. Fractions containing the SARS-CoV-2 M^pro^ were concentrated and buffer was exchanged to buffer C (20 mM Tris, 200 mM NaCl, 1 mM DTT, 1 mM EDTA, pH 7.6) using Amicon centrifugation filters. Protein was further purified by size-exclusion chromatography using an S200 column and buffer C as running buffer. Fractions with pure protein were concentrated and mixed with 50% sterile glycerol, aliquots at 4 mg/ml (~110 μM) were flash frozen in liquid nitrogen and stored at −80 °C until further use.

### Evaluation of assay buffer composition

Buffering compounds, salts, additives, polyols and detergents were evaluated by preparing a buffer solution with corresponding concentrations of the components. In the experiment evaluating the buffer components, 50 mM Tris-HCl, phosphate and HEPES buffer were prepared without the addition of other components at pH 7.5. In the experiment with salts and additives, 50 mM Tris-HCl buffer pH 7.4, was used with the addition of NaCl (in concentration 50 mM, 100 mM or 150 mM), 1 mM DTT, 1 mM TCEP or 1 mM EDTA. For the evaluation of polyols, to the 50 mM Tris-HCl buffer, pH 7.4, were added ethylene glycol (10%, 20% or 30% v/v) or glycerol (10%, 20% or 30% v/v). The influence of detergents was evaluated using 50 mM Tris-HCl buffer pH 7.4, with the addition of 0.01% detergent. In all experiments, the M^pro^ prediluted solution (final concentration 500 nM) was pipetted in the wells, corresponding buffer solution was added and the measurements were initiated by the addition of the substrate **4** (final concentration 50 μM; prepared from 10 mM DMSO stock solution). The final volume was 100 μL per well. The enzymatic activity was monitored for 15 min and determined as a slope of relative fluorescence units per second (RFU/s) for each assay additive and component. The results are expressed relative to the Tris-HCl buffer (without other components), as a mean of the triplicates and respective standard deviation. In each measurement, a corresponding solution without the addition of the protease was used as a negative control.

### Evaluation of pH value

The assay buffer (50 mM Tris-HCl, 100 mM NaCl, ethylene glycol (20% v/v) and 0.0016% Brij 58) was prepared with the pH value varying by 0.2 units in the range 7.0 - 8.0. The M^pro^ prediluted solution (final concentration 500 nM) was pipetted in the wells, corresponding buffer solution was added and the measurements were initiated by the addition of the substrate **4** (final concentration 50 μM; prepared from 10 mM DMSO stock solution). The final volume was 100 μL per well. The enzymatic activity was monitored for 15 min and determined as a slope of relative fluorescence units per second (RFU/s) for each pH value. The results are expressed relative to the assay buffer pH 7.0, as a mean of the triplicates and respective standard deviation. In each measurement, a corresponding solution without the addition of the protease was used as negative control.

### Evaluation of protease concentration

The M^pro^ prediluted solution (final concentration 0 nM, 50 nM, 100 nM, 200 nM, 300 nM or 400 nM) was pipetted in the wells, assay buffer (50 mM Tris-HCl pH 7.4, 100 mM NaCl, ethylene glycol (20% v/v) and 0.0016% Brij 58) was added and the measurements were initiated by the addition of the substrate **4** (final concentration 50 μM: prepared from 10 mM DMSO stock solution). The final volume was 100 μl per well. The enzymatic activity was monitored for 15 min and determined as a slope of relative fluorescence units per second (RFU/s) for each enzyme concentration. The results are expressed as RFU/s, as a mean of the triplicates and respective standard deviation.

### Evaluation of substrates

For the substrate screening, the M^pro^ prediluted solution (final concentration 300 nM) was pipetted in the wells, assay buffer (50 mM Tris-HCl pH 7.4, 100 mM NaCl, ethylene glycol (20% v/v) and 0.0016% Brij 58) was added and the measurements were initiated by the addition of the corresponding substrates (final concentration 50 μM: prepared from 10 mM DMSO stock solution). The final volume was 100 μl per well. The enzymatic activity was monitored for 15 min and determined as a slope of relative fluorescence units per second (RFU/s) for each substrate. The results are expressed as RFU/s, as a mean of the triplicates and respective standard deviation. In each measurement, a corresponding solution without the addition of the protease was used as a negative control.

### Kinetic measurements

Selected substrates were pipetted in the wells in the concentration range 0 - 400 mM (prepared from 10 mM DMSO stock solution), assay buffer (50 mM Tris-HCl pH 7.4, 100 mM NaCl, ethylene glycol (20% v/v) and 0.0016% Brij 58) was added and the measurements were initiated by the addition of the M^pro^ (final concentration 300 nM). The final volume was 100 μl per well. The enzymatic activity was monitored for 15 min and determined for each concentration as a slope of relative fluorescence units per second (RFU/s). The obtained values were multiplied with the correction factor for each concentration (Figures S2 and S3, Tables S1-S4). For the calculations, the enzymatic activity was also expressed as μM/s. The mean and the standard deviation of the triplicates plotted against the corresponding concentration were used to calculate the *K*_m_ and *V*_max_ values in Prism 6.01 (Graphpad Software, Inc.) using the Michaelis-Menten fit.

### Z’ value determination

The M^pro^ prediluted solution (final concentration 300 nM) together with the assay buffer (50 mM Tris-HCl pH 7.4, 100 mM NaCl, ethylene glycol (20% v/v) and 0.0016% Brij 58) was pipetted in 47 wells. The same volume of the assay buffer without the protease was pipetted in 48 wells. After 15 min preincubation, substrate **1** (final concentration 25 μM; prepared from 10 mM DMSO stock solution) was added to all wells. The final volume was 100 μL per well. The enzymatic activity was monitored for 15 min and determined as a slope of relative fluorescence units per second (RFU/s). Z’ SW and AVR values were calculated as described in literature.^39–40^

### SARS-CoV-2 Main Protease Relative Inhibition Assay

The stock solutions of the reference compounds (10 mM in DMSO) were diluted to a final concentration of 50 μM in triplicates, and preincubated for 15 min with the SARS-CoV-2 M^pro^ (300 nM) in the assay buffer (50 mM Tris-HCl pH 7.4, 100 mM NaCl, ethylene glycol (20% v/v) and 0.0016% Brij 58). The reaction was then initiated by the addition of the FRET substrate **1** (final concentration 25 μM) to obtain a final assay volume of 100 μL per well. The enzymatic activity was monitored for 15 min and determined as a slope of relative fluorescence units per second (RFU/s) for each compound. Percentage inhibition was calculated relative to a positive control (without the inhibitor), as a mean of the triplicates and respective standard deviation.

#### SARS-CoV-2 M^pro^ Assay for IC_50_ Determination

Eight inhibitor concentrations covering the range 0 – 10 or 0 – 200 μM were chosen for analysis. 1:2 serial dilutions were made starting from 10 mM stock solutions in DMSO and final concentrations were measured in triplicates. The inhibitors were preincubated for 15 min with the SARS-CoV-2 main protease (300 nM) in the assay buffer (50 mM Tris-HCl pH 7.4, 100 mM NaCl, ethylene glycol (20% v/v) and 0016% Brij 58). The enzymatic reaction was initiated by the addition of the FRET substrate **1** (final concentration 25 μM) to obtain a final assay volume of 100 μL per well. The enzymatic activity was monitored for 15 min and determined as a slope of relative fluorescence units per second (RFU/s) for each concentration. The mean and the standard deviation of the triplicates plotted against the corresponding concentration were used to determine the IC_50_-values in Prism 6.01 (Graphpad Software, Inc.) using non-linear dose-response curves with variable slopes.

## ACKNOWLEDGMENTS

We thank Malte Hermes, Max Schmitt, Katharina Eckstein and Aline Schollköpf for their involvement in SARS-CoV-2 M^pro^ production. We also thank Tobias Haupt, Mathias Büchel and Marie Hermann for preparing substrates **2, 5** and **6**, respectively, Heiko Rudy for measuring ESI high resolution spectra and Natascha Stefan for technical assistance.

## ABBREVIATIONS USED

CHAPS: 3-[(3-cholamidopropyl)dimethylammonio]-l-propanesulfonate
Dap(Dnp): *N*^β^-2,4-dinitrophenyl-L-diaminopropionic acid
DENV: dengue vims
DTT: dithiothreitol
EDTA: ethylenediaminetetraacetic acid
EG: ethylene glycol
Fmoc: fluorenylmethyloxycarbonyl protecting group
FRET: Forster resonance energy transfer
Gly: glycerol
HEPES: 2-[4-(2-hydroxyethyl)piperazin-l-yl]ethanesulfonic acid
MERS: Middle East respiratory syndrome
M^pro^: SARS coronavirus main proteinase
RFU: relative fluorescence units
SARS: severe acute respiratory syndrome
TCEP: tris(2-carboxyethyl)phosphine
Tris: 2-amino-2-(hydroxymethyl)propane-l,3-diol
WNV: West Nile virus

