## Supporting Information for "Efficiency Improvements and Discovery of New Substrates for a SARS-CoV-2 Main Protease FRET Assay"

#### Measurement of the activity of C144A mutant protease

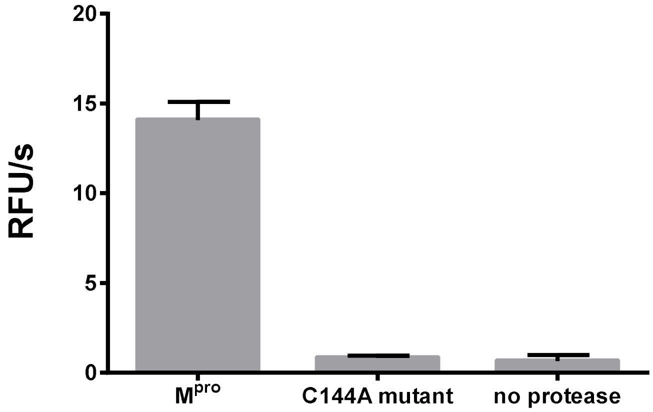

**Figure S1.** The measurement was performed in Tris buffer, pH 7.6 at the enzyme concentration 1 µM and with substrate 3 at concentration 50 µM.

**Measurement 2-Abz calibration curve**

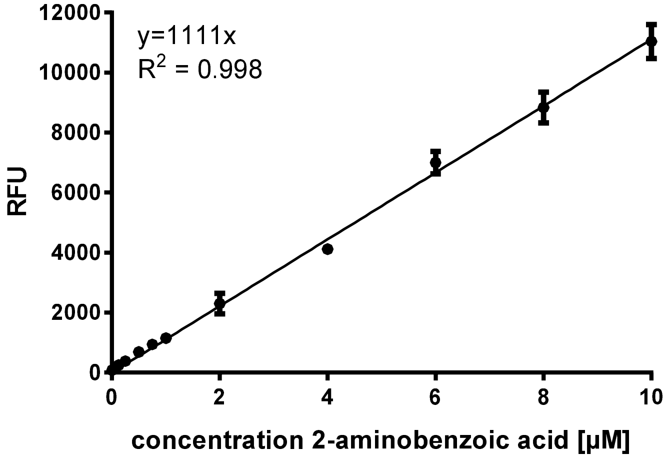

**Figure S2.** Linear 2-aminobenzoic acid calibration curve.

### Determination of FRET substrate correction factors

Correction factors were determined for each substrate as described previously.[1-2]

**Table S1. Correction factors for substrate 1**

| Substrate concentration [µM] | Correction factors |
|---|---|
| 400 | 0,4585 |
| 300 | 0,5415 |
| 200 | 0,7020 |
| 150 | 0,7541 |
| 100 | 0,7970 |
| 50 | 0,9170 |
| 25 | 0,9639 |
| 12,5 | 1,0003 |
| 0 | 1,0000 |

**Table S2. Correction factors for substrate 2**

| Substrate concentration [µM] | Correction factors |
|---|---|
| 400 | 0,4523 |
| 300 | 0,6108 |
| 200 | 0,7103 |
| 150 | 0,7916 |
| 100 | 0,8596 |
| 50 | 0,9294 |
| 25 | 1,0122 |
| 12,5 | 1,0189 |
| 0 | 1,0000 |

**Table S3. Correction factors for substrate 3**

| Substrate concentration [µM] | Correction factors |
|---|---|
| 400 | 0,6383 |
| 300 | 0,7908 |
| 200 | 0,8846 |
| 150 | 0,9674 |
| 100 | 1,1174 |
| 50 | 1,0940 |
| 25 | 1,0840 |
| 12,5 | 1,0840 |
| 0 | 1,0000 |

**Table S4. Correction factors for substrate 4**

| Substrate concentration [µM] | Correction factors |
|---|---|
| 400 | 0,3746 |
| 300 | 0,4508 |
| 200 | 0,6184 |
| 150 | 0,7029 |
| 100 | 0,7835 |
| 50 | 0,9552 |
| 25 | 0,9877 |
| 12,5 | 0,9944 |
| 0 | 1,0000 |

**Michaelis–Menten curves**

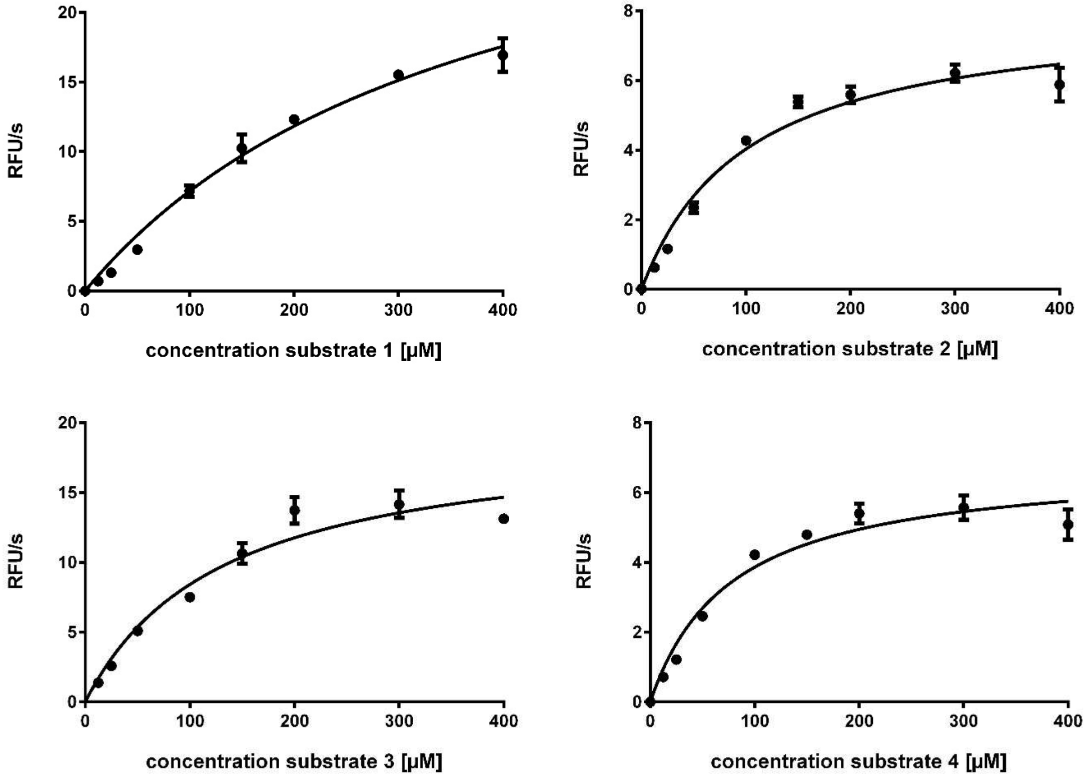

**Figure S3.** Michaelis–Menten curves of the interaction of FRET substrates with the SARS-CoV-2 $M^{pro}$. Substrate cleavage velocity in RFU/s. All curves were corrected for inner filter effects.[2] All measurements from triplicate data.

**Reagents and Solvents.** All chemicals for the synthesis were obtained from Sigma-Aldrich (Germany), Alfa Aesar (Germany), TCI Europe (Belgium) and were of analytical grade. Solvents were used as obtained from the commercial suppliers.

**Equipment and Analytical Methods**. NMR spectra were recorded on Varian NMR instrument at 300 MHz, 300 K; Chemical shifts (δ) are given in parts per million (ppm). Residual peaks of nondeuterated solvents were used as internal standard: DMSO-$d_6$ (δ ppm: 2.50). Coupling constants (*J*) are given in hertz (Hz). Multiplicity is reported as s (singlet), d (doublet), t (triplet), dd (doublet of doublet), m (multiplet). Mass spectra (HR-ESI) of all compounds were measured on a Bruker micrOTOF-Q II instrument. Analysis of the compounds and intermediates was carried out using solutions in water, methanol, acetonitrile or water/methanol.

**Synthesis of substrates 1 – 8**

The substrates were synthesized using the solid phase peptide synthesis. The reactions were performed in 5 mL syringes with shaking at each coupling and cleavage step. The α-amino groups in all amino acids were protected with Fmoc protecting group, whereas the reactive groups in the side chains of respective amino acids were protected with appropriate protecting groups. The synthesis was performed on CTC resin (100 mg, 1.6 mmol/g). At the beginning, the resin was swelled with dichloromethane for minimally 30 minutes and afterwards, washed with DMF (3 × 1 mL). For the coupling of the first amino acid, the amino acid (1.0 equiv) was dissolved in DMF (2 mL) and NMM (4.0 equiv) was added. The resulting solution was drawn into the syringe and the reaction proceeded for at least 2.5 h with shaking. The resin was washed with DMF (3 × 1 mL), dichloromethane (3 × 1 mL) and again with DMF (3 × 1 mL). After the first coupling step, the resin was capped with a mixture of dichloromethane : methanol : NMM in ratio 80 : 15 : 5 (2 mL). The capping proceeded two times for 30 minutes. The resin was washed with DMF (3 × 1 mL), dichloromethane (3 × 1 mL) and again with DMF (3 × 1 mL). Fmoc group was then cleaved using 25% piperidine in DMF (2 mL) for 10 minutes with shaking. The cleavage step was repeated two times. Before the next coupling step, the resin was washed with DMF (6 × 1 mL), dichloromethane (3 × 1 mL) and again with DMF (3 × 1 mL). The second amino acid (1.0 equiv) and HATU (3.0 equiv) were dissolved in DMF (2 mL), and NMM (4.0 equiv) was added. The resulting solution was drawn into the syringe and the reaction proceeded for at least 2.5 h with shaking. The resin was washed with DMF (3 × 1 mL), dichloromethane (3 × 1 mL) and again with DMF (3 × 1 mL). The Fmoc group was cleaved using 25% piperidine in DMF (2 mL) as described above and the resin was washed with DMF (6 × 1 mL), dichloromethane (3 × 1 mL) and again with DMF (3 × 1 mL). The consecutive amino acids and the 2-Abz cap were coupled following the same procedure. After the last cleavage step, the resin was washed with diethyl ether (5 × 1 mL) and dried overnight under reduced pressure. The peptide was cleaved from the resin using a mixture trifluoroacetic acid : water: TIPS in ratio 95 : 2.5 : 2.5. The mixture (2 mL) was drawn into the syringe and the reaction proceeded for 2 h with shaking. The content of the syringe was released into a Falcon tube containing an ice-cold diethyl ether (35 mL). The cleavage was repeated one time. The Falcon tube was centrifuged at 4000 RCF for 5 minutes at 4 °C, after which the supernatant was removed. The pellet was subjected to preparative HPLC purification to obtain the pure substrates. The conditions of the purification were as follows: RP-18 pre- and main column (Reprospher 100 C-18-DE, Dr. Maisch GmbH, Germany, 5 µm, precolumn 30 × 16 mm, main column 125 × 16 mm); eluent A, water (0.1% TFA); eluent B, methanol (0.1% TFA) or acetonitrile (0.1% TFA); 0–2.5 min, 10% B; 2.6–23.5 min, gradient 10% B to 100% B; 23.6–26.0 min, 100% B; 26.1–30.0 min, 10% B; flow rate, 8 mL/min; $\lambda$ = 214, 254, and 280 nm.

Substrates **7** and **8** with the sequences Abz-Nle-Lys-Arg-Arg-Ser-3-($NO_2$)Tyr-$NH_2$ and 2-Abz-Gly-Lys-Lys-Arg-Gly-Tyr(3-$NO_2$)-Ala-Lys-$NH_2$, respectively, were synthesized by solid phase peptide synthesis on a Rink amide resin using N-terminal Fmoc protected amino acids, HATU as a coupling reagent and DIPEA as a base, according to the previously established procedure.[1]

**Substrate 1.** Obtained as yellow powder (31 mg, 15%). Purity > 99%, determined by HPLC; retention time 2.7 min. HRMS (ESI): m/z $[M + H]^+$ calcd for $C_{49}H_{74}N_{15}O_{17}$, 1144.5382; found, 1144.5367.

**Substrate 2.** Obtained as yellow powder (21 mg, 10%). Purity > 99%, determined by HPLC; retention time 2.6 min min. HRMS (ESI): m/z $[M + H]^+$ calcd for $C_{50}H_{76}N_{15}O_{18}$, 1174.5487; found, 1174.5524.

**Substrate 3.** Obtained as yellow powder (12 mg, 6%). Purity > 99%, determined by HPLC; retention time 2.8 min. HRMS (ESI): m/z $[M + H]^+$ calcd for $C_{52}H_{80}N_{15}O_{17}$, 1186.5851; found, 1186.5837.

**Substrate 4.** Obtained as yellow powder (38 mg, 18%). Purity 99%, determined by HPLC; retention time 3.1 min. HRMS (ESI): m/z $[M + H]^+$ calcd for $C_{52}H_{80}N_{17}O_{18}$, 1230.5862; found, 1230.5818.

**Substrate 5.** Obtained as yellow powder (46 mg, 22%). Purity 95%, determined by HPLC; retention time 2.4 min. HRMS (ESI): m/z $[M + H]^+$ calcd for $C_{51}H_{75}N_{16}O_{17}$, 1183.5491; found, 1183.5491.

**Substrate 6.** Obtained as yellow powder (14 mg, 9%). Purity 90%, determined by HPLC; retention time 2.8 min. HRMS (ESI): m/z $[M + H]^+$ calcd for $C_{37}H_{52}N_9O_{13}$, 830.3679; found, 830.3700.

**Substrate 7.** Obtained as yellow powder (108 mg, 65%). Purity > 99%, determined by HPLC; retention time 3.1 min. HRMS (ESI): m/z $[M + H]^+$ calcd for $C_{43}H_{69}N_{16}O_{11}$, 985.5326; found, 985.6310.

**Substrate 8.** Obtained as yellow powder (84 mg, 23%). Purity 97%, determined by HPLC; retention time 1.7 min. HRMS (ESI): m/z $[M + H]^+$ calcd for $C_{47}H_{76}N_{17}O_{12}$, 1070.5854; found, 1070.5870.

### Synthesis of inhibitors

*General Procedure for Synthesis of Ester Compounds*

Ester compounds were synthesized in analogy to previously described procedures.[3-4] In short, EDCl · HCl (1.0 equiv), HOBt (1.0 equiv) and the hydroxyl derivative (1.0 equiv) were given to a solution of the carboxylic acid (1.0 equiv) in DMF (3 mL) at room temperature. DIPEA (2.0 equiv) was added and the solution was stirred overnight. The mixture was dissolved in ethyl acetate and washed 4 times with $H_2O$. The organic layer was dried over anhydrous MgSO4 and concentrated *in vacuo*. The compounds were purified by preparative RP-HPLC on an ÄKTA Purifier, GE Healthcare (Germany), with an RP-18 pre and main column (Rephospher, Dr. Maisch GmbH, Germany, C18-DE, 5 µm, 30 mm × 16 mm and 120 mm × 16 mm). The following conditions were used: eluent A, water (0.1% TFA); eluent B, methanol (0.1% TFA) or eluent A, water (0.1% TFA); eluent B, acetonitrile (0.1% TFA); flow rate, 8 mL/min; and gradient, 10% B (2.5 min), 100% B (23.5 min), 100% B (26 min), 10% B (26.1 min), and 10% B (30 min). Detection was performed at 214, 254, and 280 nm. After purification, the organic solvent was evaporated, and the compounds and intermediates were freeze-dried in $H_2O$/ACN and stored at -20 °C.

**5-Chloropyridin-3-yl thiophene-2-carboxylate (MAC-5576)**

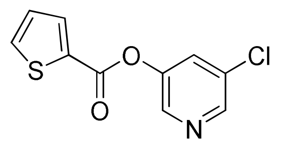

MAC-5576 was synthesized according to general procedure for synthesis of ester compounds from thiophene-2-carboxylic acid (35 mg, 0.27 mmol), EDCl · HCl (52 mg, 0.27 mmol), HOBt (37 mg, 0.27 mmol), 5-chloro-3-pyridinol (35 mg, 0.27 mmol) and DIPEA (0.093 mL, 0.55 mmol). Obtained as a white powder (61 mg, 94% yield). $^1$H NMR (300 MHz, DMSO-d$_6$) δ 8.87–8.38 (m, 2H), 8.15 (dd, *J* = 2.5, 1.1 Hz, 1H), 8.12 (s, 1H), 7.64 (dd, *J* = 3.6, 0.7 Hz, 1H), 6.83 (dd, *J* = 3.6, 1.7 Hz, 1H). $^{13}$C NMR (APT, 75 MHz, DMSO-d$_6$) δ 159.5, 145.9, 142.1, 136.2, 136.1, 135.9, 135.9, 130.8, 130.2, 128.8. HRMS (ESI): *m/z* [M + H]$^+$ calcd for $C_{10}H_7ClNO_2S$: 239.9881, found: 239.9877.

**5-chloropyridin-3-yl furan-2-carboxylate (FE-1)**

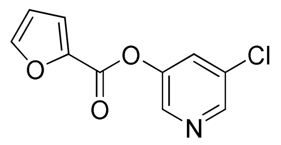

**FE-1** was synthesized according to general procedure for synthesis of ester compounds from furan-2-carboxylic acid (31 mg, 0.28 mmol), EDCl · HCl (53 mg, 0.28 mmol), HOBt (37 mg, 0.28 mmol), 5-chloro-3-pyridinol (36 mg, 0.28 mmol) and DIPEA (0.094 mL, 0.55 mmol). Obtained as a white powder (58 mg, 94% yield). $^1$H NMR (300 MHz, DMSO-d$_6$) δ 8.76–8.50 (m, 2H), 8.15 (dd, *J* = 5.0, 1.1 Hz, 1H), 8.13 (s, 1H), 8.07 (dd, *J* = 3.8, 1.2 Hz, 1H), 7.33 (dd, *J* = 4.9, 3.9 Hz, 1H). $^{13}$C NMR (APT, 75 MHz, DMSO-d$_6$) δ 155.6, 149.2, 146.0, 145.9, 142.2, 142.1, 130.2, 130.2, 121.1, 113.0. HRMS (ESI): *m/z* [M + H]$^+$ calcd for $C_{10}H_7ClNO_3$: 224.0109, found: 224.0103.

### HPLC Purity of FRET Substrates and Reference Compounds

Purity of inhibitors was determined by HPLC on a Jasco HPLC system with a Jasco UV-2070 Plus Intelligent UV/VIS Detector on an RP-18 column (ReproSil-Pur-ODS-3, Dr. Maisch GmbH, Germany, 5 µm, 50 mm × 2 mm) using the following method: eluent A: water (0.1% TFA); eluent B: acetonitrile (0.1% TFA); injection volume: 10 µL; flow rate: 1 mL/min; and gradient: 1% B (0.2 min), 100% B (7 min), 100% B (8 min), 1% B (8.1 min), and 1% B (10 min). Chromatograms recorded at 254 nm were used for purity assessment.

**Table S5. RP-HPLC data of substrates and compounds**

| Compound | HPLC $t_R$ (min) | HPLC purity (254 nm) |
|---|---|---|
| **Substrate 1** | 2.7 | > 99% |
| **Substrate 2** | 2.6 | > 99% |
| **Substrate 3** | 2.8 | > 99% |
| **Substrate 4** | 3.1 | 99% |
| **Substrate 5** | 2.4 | 95% |
| **Substrate 6** | 2.8 | 90% |
| **Substrate 7** | 3.1 | >99% |
| **Substrate 8** | 1.7 | 97% |
| **MAC-5576** | 3.2 | > 95% |
| **FE-1** | 2.7 | > 95% |

**1H und 13C NMR Spectra**

Compound **MAC-5576**, $^{1}$H NMR (300 MHz, DMSO-$d_6$)

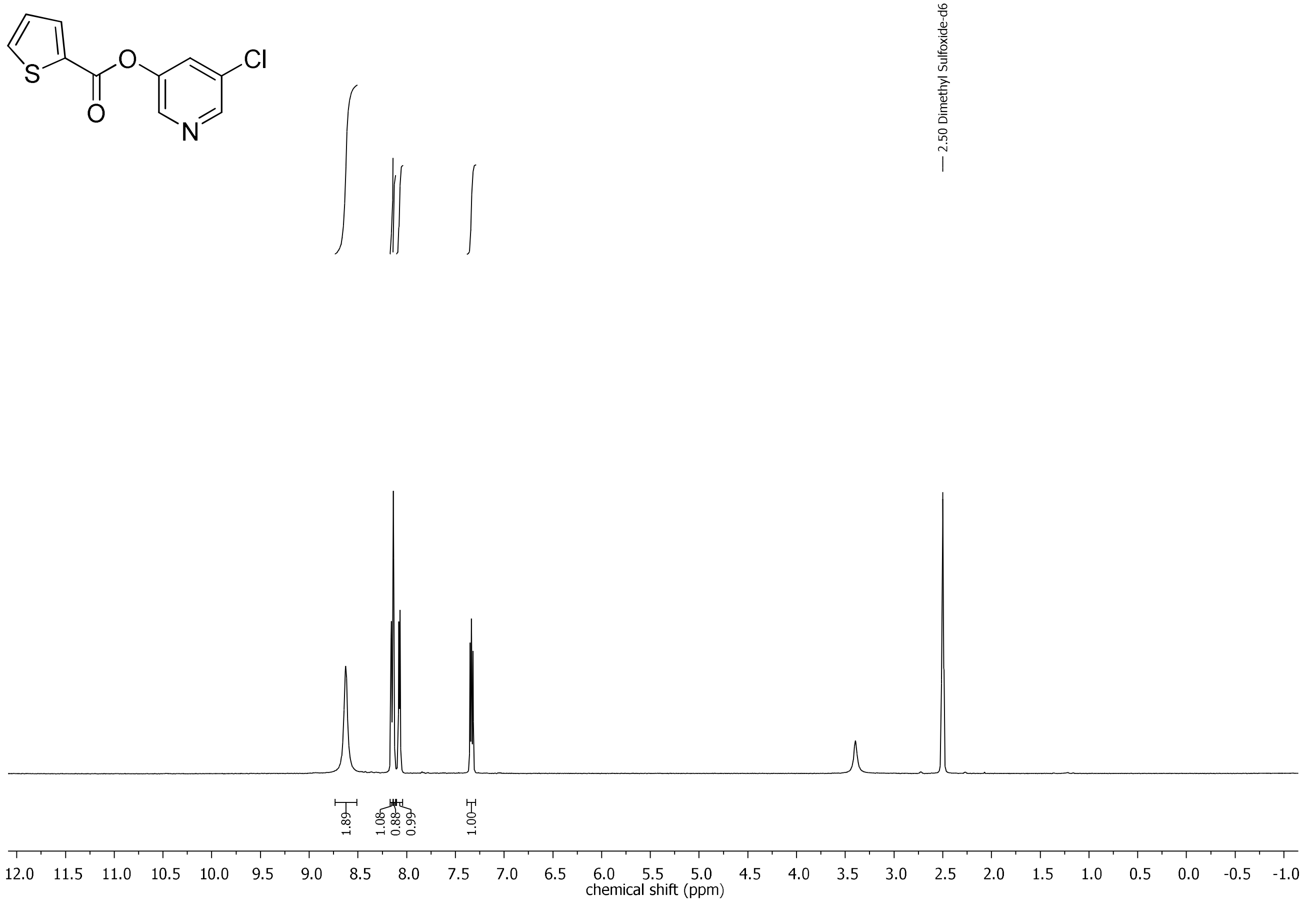

Compound **MAC-5576**, $^{13}C$ NMR (APT, 75 MHz, DMSO-$d_6$)

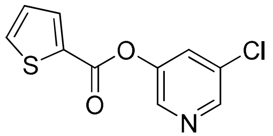

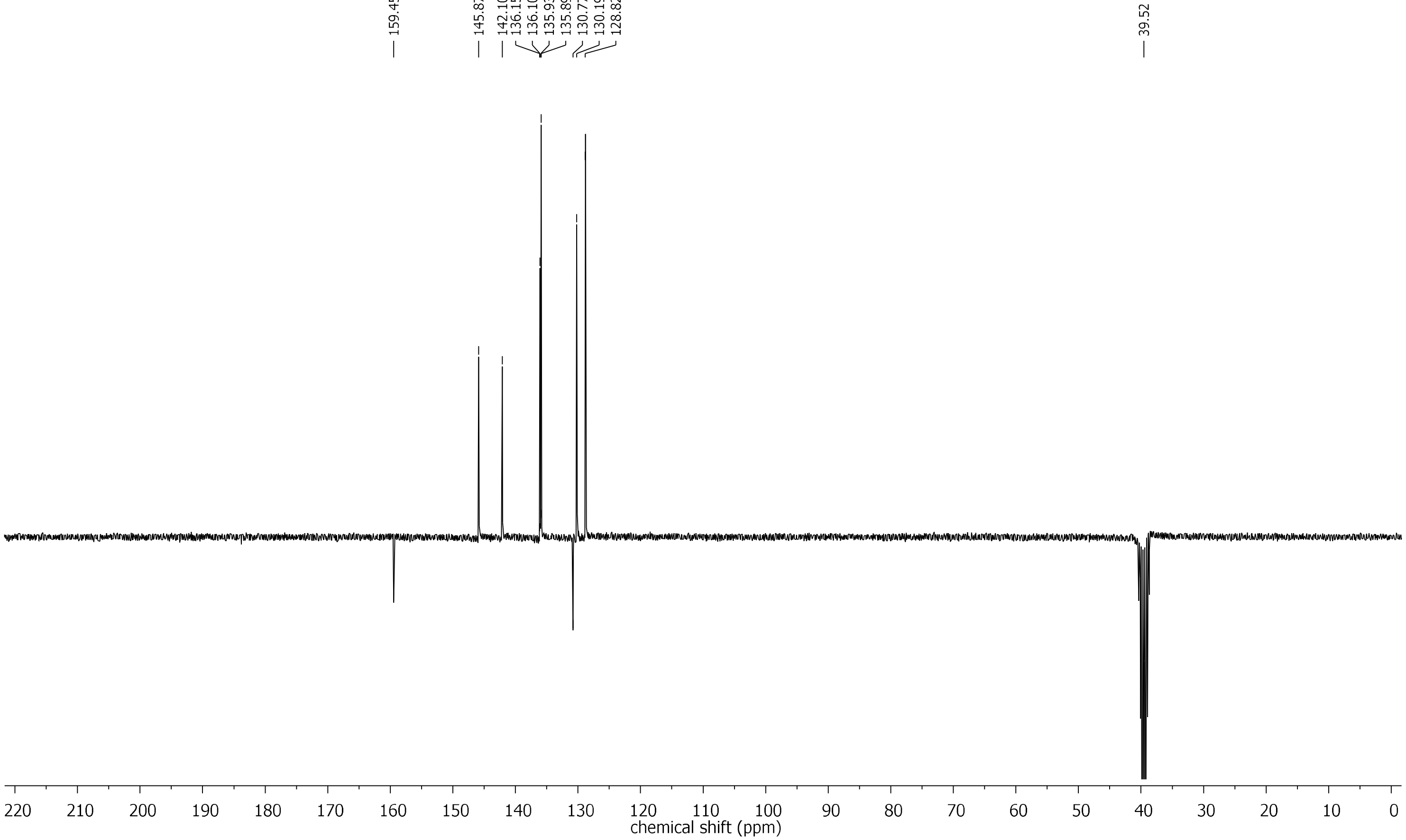

Compound **FE-1**, $^1$H NMR (300 MHz, DMSO-$d_6$)

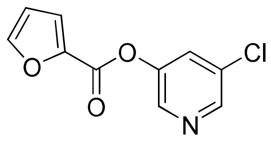

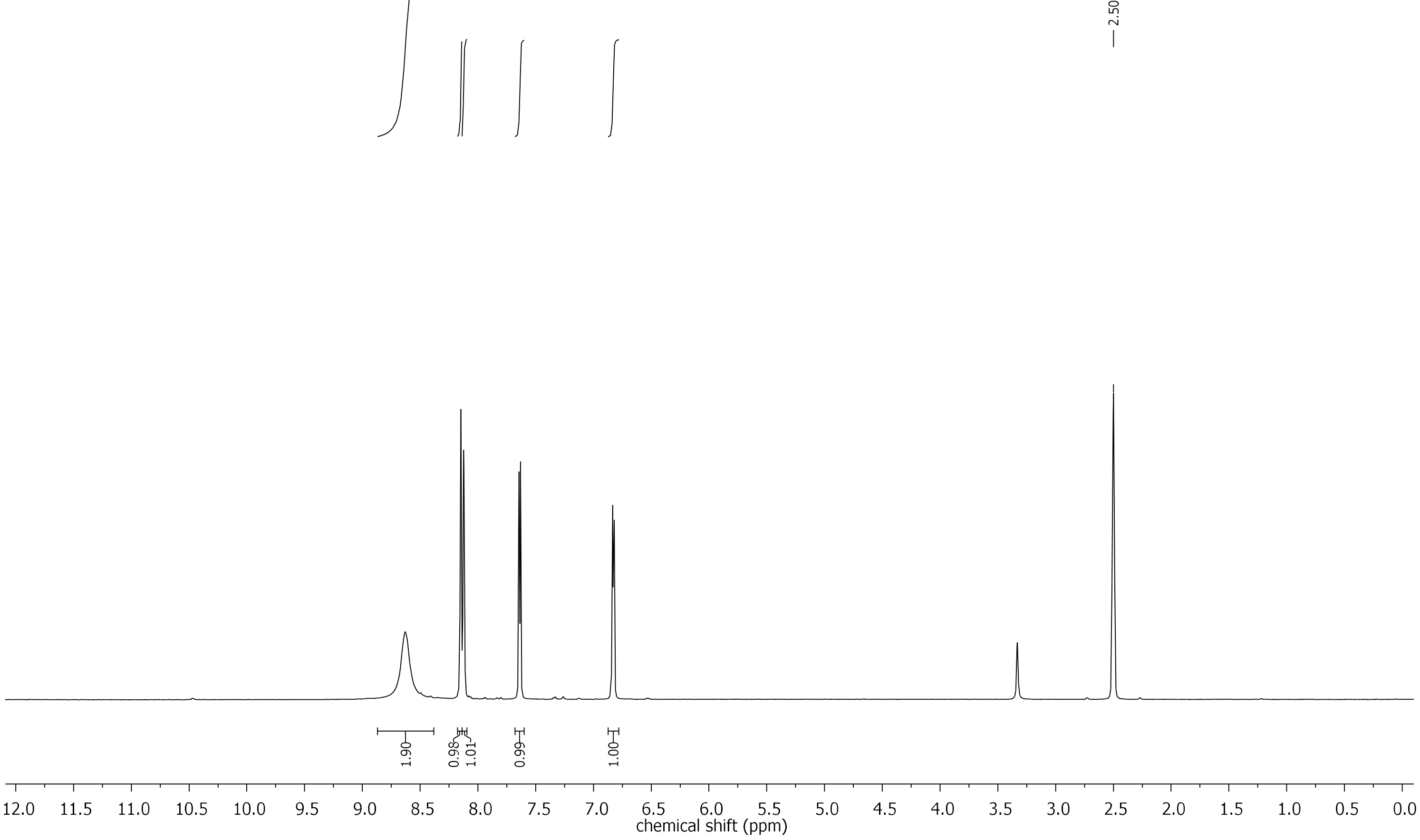

Compound **FE-1**, $^{13}$C NMR (APT, 75 MHz, DMSO-d$_6$)

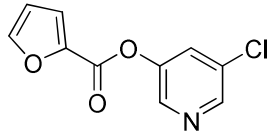

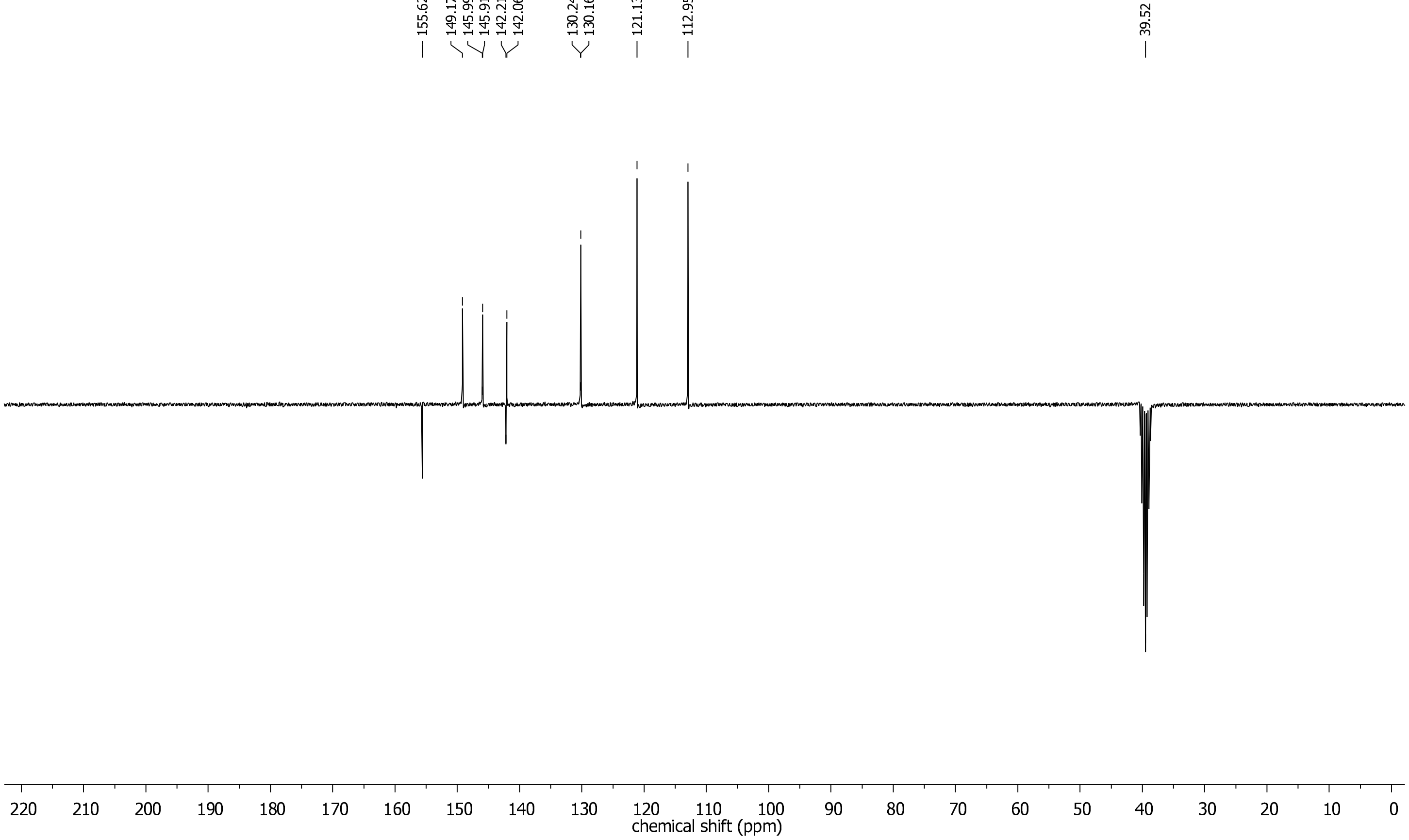

### HPLC analysis of the substrate purity

#### Substrate 1

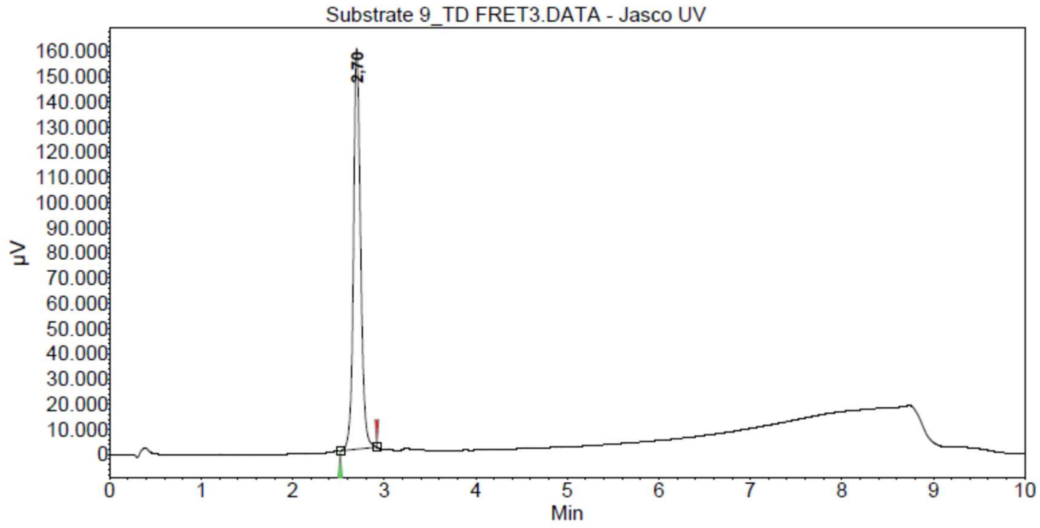

**Peak results :**

| Index | Name | Time [Min] | Quantity [% Area] | Height [µV] | Area [µV.Min] | Area % [%] |
|---|---|---|---|---|---|---|
| 1 | UNKNOWN | 2,70 | 100,00 | 159204,9 | 14283,0 | 100,000 |
| Total | | | 100,00 | 159204,9 | 14283,0 | 100,000 |

#### Substrate 2

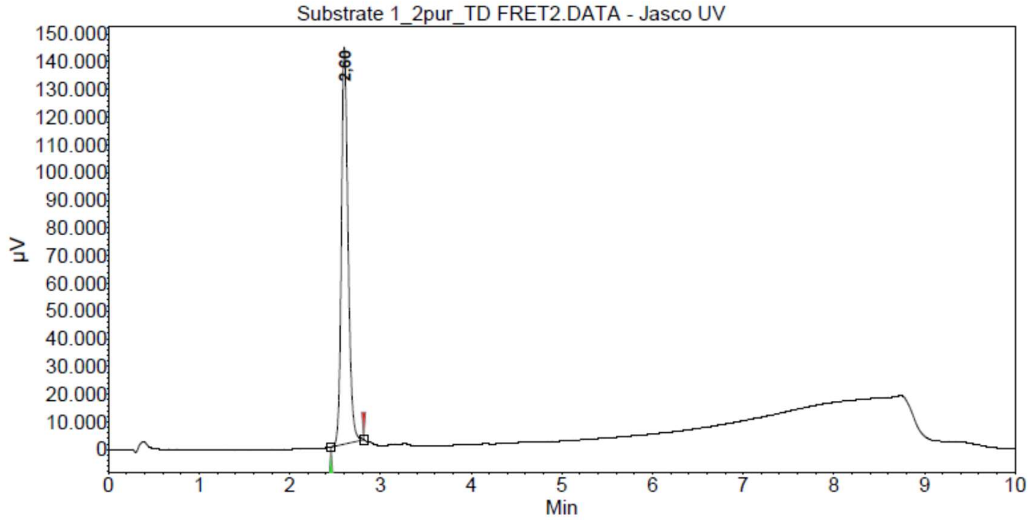

**Peak results :**

| Index | Name | Time [Min] | Quantity [% Area] | Height [µV] | Area [µV.Min] | Area % [%] |
|---|---|---|---|---|---|---|
| 1 | UNKNOWN | 2,60 | 100,00 | 143417,7 | 12250,8 | 100,000 |
| Total | | | 100,00 | 143417,7 | 12250,8 | 100,000 |

**Substrate 3**

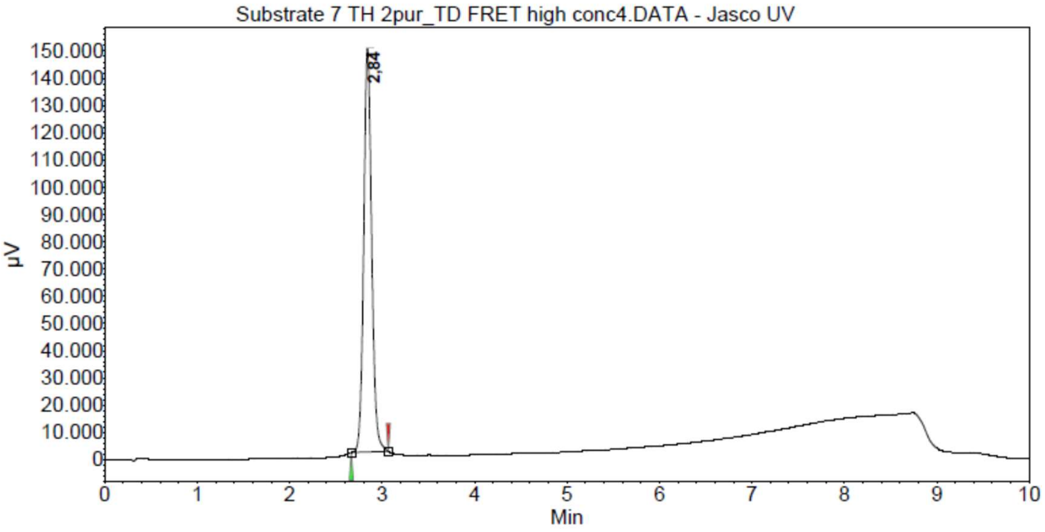

**Peak results :**

| Index | Name | Time [Min] | Quantity [% Area] | Height [µV] | Area [µV.Min] | Area % [%] |
|---|---|---|---|---|---|---|
| 1 | UNKNOWN | 2,84 | 100,00 | 148278,4 | 14753,6 | 100,000 |
| Total | | | 100,00 | 148278,4 | 14753,6 | 100,000 |

**Substrate 4**

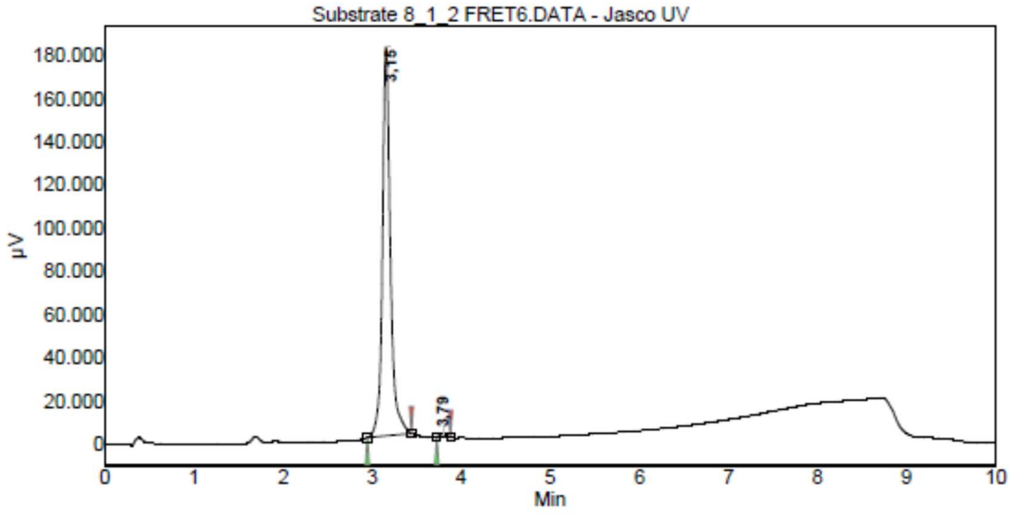

**Peak results :**

| Index | Name | Time [Min] | Quantity [% Area] | Height [µV] | Area [µV.Min] | Area % [%] |
|---|---|---|---|---|---|---|
| 1 | UNKNOWN | 3,15 | 99,26 | 180154,8 | 18596,1 | 99,260 |
| 2 | UNKNOWN | 3,79 | 0,74 | 1695,3 | 138,7 | 0,740 |
| Total | | | 100,00 | 181850,1 | 18734,8 | 100,000 |

### Substrate 5

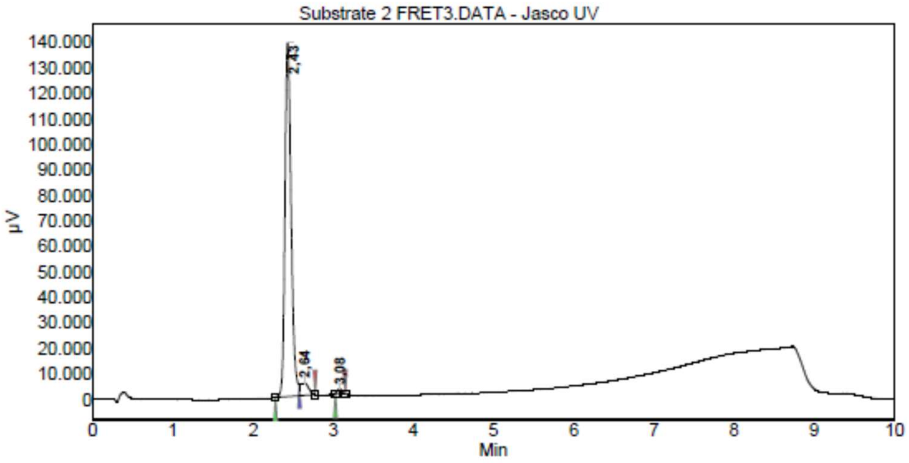

**Peak results :**

| Index | Name | Time [Min] | Quantity [% Area] | Height [µV] | Area [µV.Min] | Area % [%] |
|---|---|---|---|---|---|---|
| 1 | UNKNOWN | 2,43 | 94,58 | 138785,6 | 12385,1 | 94,577 |
| 2 | UNKNOWN | 2,64 | 4,50 | 4756,0 | 589,0 | 4,498 |
| 3 | UNKNOWN | 3,08 | 0,93 | 1782,2 | 121,2 | 0,925 |
| Total | | | 100,00 | 145323,8 | 13095,3 | 100,000 |

### Substrate 6

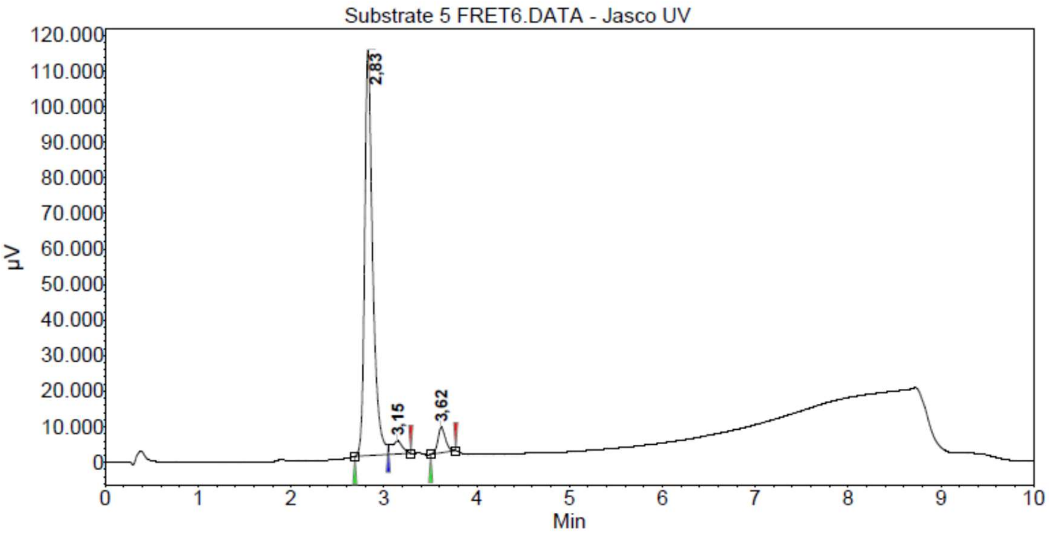

**Peak results :**

| Index | Name | Time [Min] | Quantity [% Area] | Height [µV] | Area [µV.Min] | Area % [%] |
|---|---|---|---|---|---|---|
| 2 | UNKNOWN | 2,83 | 90,49 | 114158,4 | 11401,6 | 90,492 |
| 3 | UNKNOWN | 3,15 | 4,15 | 3718,2 | 523,4 | 4,154 |
| 1 | UNKNOWN | 3,62 | 5,35 | 7142,2 | 674,5 | 5,354 |
| Total | | | 100,00 | 125018,7 | 12599,5 | 100,000 |

### Substrate 7

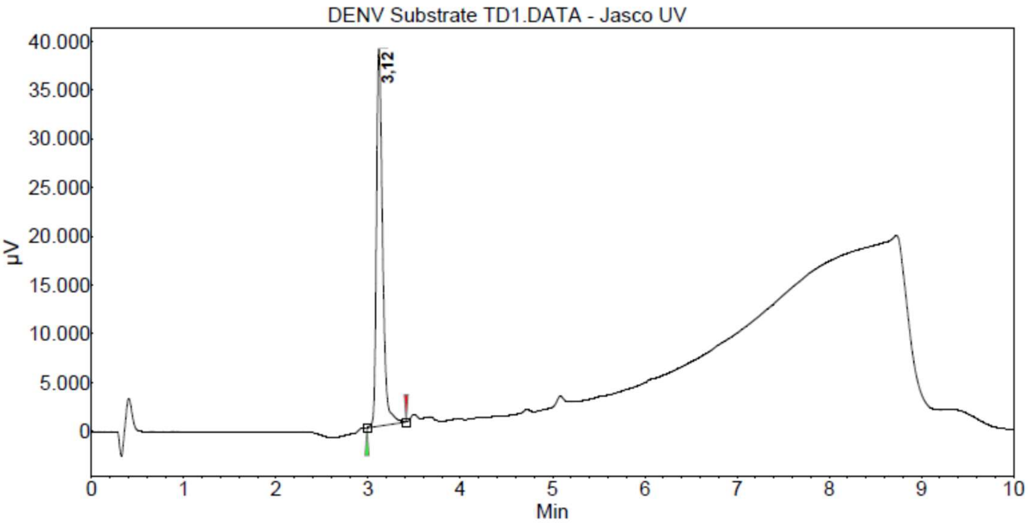

**Peak results :**

| Index | Name | Time [Min] | Quantity [% Area] | Height [μV] | Area [μV.Min] | Area % [%] |
|---|---|---|---|---|---|---|
| 1 | UNKNOWN | 3,12 | 100,00 | 38728,7 | 2976,3 | 100,000 |
| Total | | | 100,00 | 38728,7 | 2976,3 | 100,000 |

### Substrate 8

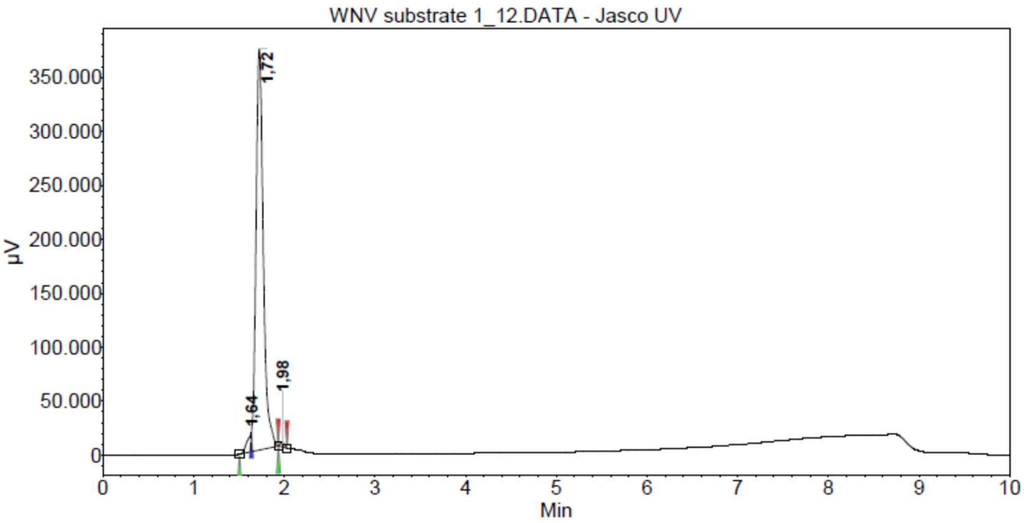

**Peak results :**

| Index | Name | Time [Min] | Quantity [% Area] | Height [μV] | Area [μV.Min] | Area % [%] |
|---|---|---|---|---|---|---|
| 1 | UNKNOWN | 1,64 | 2,56 | 18610,2 | 912,6 | 2,556 |
| 3 | UNKNOWN | 1,72 | 96,86 | 371843,1 | 34588,7 | 96,858 |
| 2 | UNKNOWN | 1,98 | 0,59 | 4095,9 | 209,6 | 0,587 |
| Total | | | 100,00 | 394549,2 | 35710,9 | 100,000 |

**DMSO**

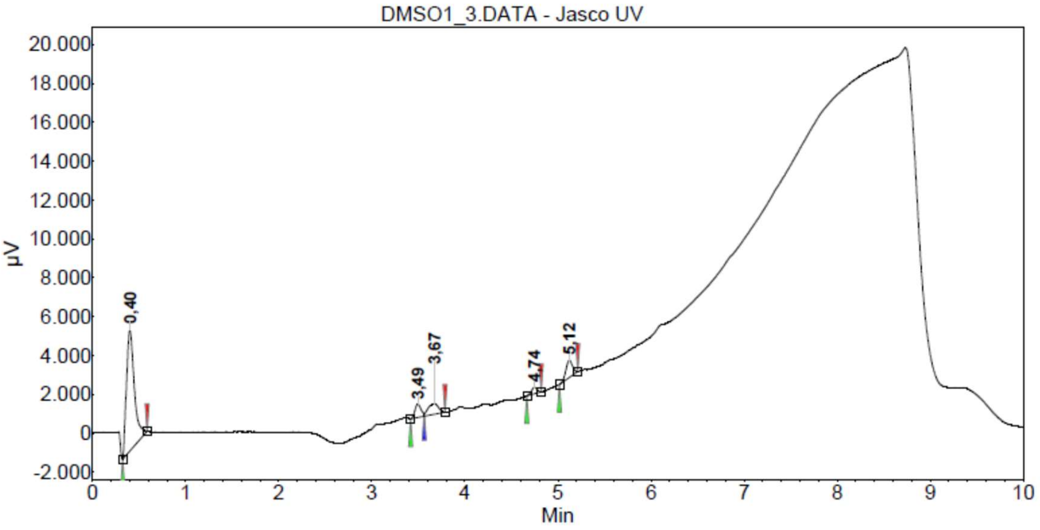

**Peak results :**

| Index | Name | Time [Min] | Quantity [% Area] | Height [µV] | Area [µV.Min] | Area % [%] |
|---|---|---|---|---|---|---|
| 1 | UNKNOWN | 0,40 | 73,76 | 6200,4 | 608,3 | 73,761 |
| 2 | UNKNOWN | 3,49 | 6,25 | 689,1 | 51,5 | 6,250 |
| 3 | UNKNOWN | 3,67 | 8,47 | 532,1 | 69,8 | 8,465 |
| 4 | UNKNOWN | 4,74 | 2,77 | 329,9 | 22,8 | 2,769 |
| 5 | UNKNOWN | 5,12 | 8,75 | 886,9 | 72,2 | 8,755 |
| Total | | | 100,00 | 8638,4 | 824,8 | 100,000 |
